# The long non-coding RNA NEAT1 regulates the transcriptional landscape of cardiomyocytes

**DOI:** 10.1101/2024.06.27.600932

**Authors:** Priyanka Pant, Prakash Sivakumar, Disha Nanda, N Sai Ram, Perundurai Dhandapany, Thomas Thum, Regalla Kumarswamy

**Affiliations:** CSIR-Center for Cellular and Molecular Biology (CCMB), Hyderabad, India; Academy of Scientific and Innovative Research, Ghaziabad (AcSIR), India; Institute of Institute for Stem Cell Science and Regenerative Medicine (inSTEM), Bengaluru; Institute of Molecular and Translational Therapeutic Strategies (IMTTS) and Rebirth Center for Translational Regenerative Therapies, Hannover Medical School, Hannover, Germany

## Abstract

Long non-coding RNAs (lncRNAs) play a crucial role in fine-tuning the gene expression. In the present study, we identified that the lncRNA NEAT1 is upregulated in failing murine hearts and in human hypertrophic cardiomyopathies. Further investigations demonstrated that NEAT1 expression is regulated by calcium dependent NFAT (nuclear factor of activated T cells). Overexpression of NEAT1 led to increased cardiomyocyte size and elevated expression of cardiac stress markers while its depletion had the opposite effect. Through transcriptomic analysis in NEAT1-KO cells, we demonstrated the cis-regulatory role of NEAT1, wherein it maintains the transcription of neighboring genes by interacting with the coactivator p300. Additionally, NEAT1 depletion resulted in decreased expression of GRK2 (G protein-coupled receptor kinase 2), a key player in the development of cardiac hypertrophy, suggesting that NEAT1 contributes to the development of cardiac hypertrophy by regulating GRK2 expression. Consistent with this, Neat1^-/-^ mice are resistant to β-adrenergic stimulation. Overall, our study provides enough evidence that NEAT1 is a calcium-regulated, pro-hypertrophic lncRNA that exerts significant control over the transcriptional landscape of cardiomyocytes during heart failure.

## Introduction

Pathological cardiac hypertrophy is a major contributor to heart failure. Numerous studies have shown that inhibiting cardiac hypertrophy under stress conditions helps preserve cardiac function. Currently, the available treatment options are mostly limited to β-blockers, angiotensin receptor-neprilysin inhibitors (ARNi), and calcium agonists.^1^ Long non-coding RNAs are transcripts over >200nts length lacking the ability to code for any protein. Several lncRNAs have been identified to contribute to the development of cardiac hypertrophy. Heart-specific lncRNA *Mhrt* promotes the expression of Myh7 through its interaction with the chromatin remodeler Brg1.^2^ *Chast* promotes cardiac hypertrophy by targeting autophagy, and depletion of *Chast* using gapmers has been shown to regress hypertrophy.^3^ Other examples of lncRNAs involved in cardiac hypertrophy include NRON,^4^ CHRF,^5^ Ahit,^6^ Chaer,^7^ DACH1,^8^ Zfas1,^9^ and H19.^10^ These findings highlight the potential of lncRNAs as therapeutic targets for cardiac diseases.

Restoring calcium homeostasis is crucial for improving cardiac contractility in failing hearts. Various preclinical studies have highlighted the efficacy of AAV-mediated SERCA2a gene therapy in reversing pathological cardiac remodeling.^11^ We conducted a microarray-based profiling of the long non-coding RNAs (lncRNAs) in a chronic post-myocardial infarction heart failure model treated with AAV9.SERCA2a gene therapy.^12^ Several lncRNAs were de regulated in failing hearts. Interestingly, expression of an evolutionarily conserved lncRNA NEAT1 was highly increased in failing hearts and its expression restored to normal levels in the animals that received SERCA2A gene therapy. Further in-vitro and in-vivo experimental studies revealed that NEAT1 regulates cardiac hypertrophy. RNA-sequencing and Stable isotope labeling by amino acids in cell culture (SILAC) experiments revealed that neighboring genes of NEAT1 are particularly affected upon NEAT1 silencing. Mechanistically, NEAT1 enhances recruitment of histone acetyl transferase p300 to its neighboring genes, including β-adrenergic receptor kinase 2 (GRK2) and facilitates their active transcription. Our findings suggest that NEAT1 can be a potential therapeutic target for cardiac hypertrophy.

## Results

### NEAT1 expression is increased during heart failure

We previously profiled miRNA expression in ventricular tissues of rats with heart failure (HF) and HF hearts that received AAV9-SERCA2a.^12^ In the present study, we profiled lncRNA expression in those samples and found that several lncRNAs were deregulated in failing hearts (Table S1, Figure 1A, B). Evolutionarily conserved lncRNA NEAT1 was highly upregulated in failing hearts and its expression was restored to normal levels in animals that received SERCA2a gene therapy. (Figure 1B). Independent qRT-PCR experiments confirmed the same (Figure 1C). In humans, NEAT1 is expressed as two isoforms, a 3.7 kb smaller isoform (NEAT1.1) and a 23 kb long isoform (NEAT1.2). Expression of both the isoforms and cardiac stress markers ANP and β-MHC increased in neonatal rat cardiomyocytes upon stimulation with isoproterenol (ISO) (Figure 1D). Similar results were obtained in rat cardiomyoblast cell line H9C2, human cardiomyocyte cell line AC16, and mouse cardiomyocyte cell line HL-1, where treatment with isoproterenol and phenylephrine (PE) led to elevated expression of NEAT1, NEAT1.2, along with ANP (Figure S1A, B, C). Cardiac Neat1 expression was tested during cardiac hypertrophy induced by pressure overload (TAC) or upon treatment with β-adrenergic receptor agonist isoproterenol. Both the models displayed cardiac hypertrophy with increased heart size, cardiomyocytes size, and increased levels of cardiac stress marker expression (Figure S2 and S3, Table S2 and Table S3). Neat1 expression increased in both these cardiac hypertrophy models as ejection fraction (EF) decreased (Figure 1E, F). Interestingly, Neat1 is highly enriched in cardiomyocyte fraction when compared with non-myocyte fraction of the heart in mice (Figure 1G and S1D).

**Figure 1.**
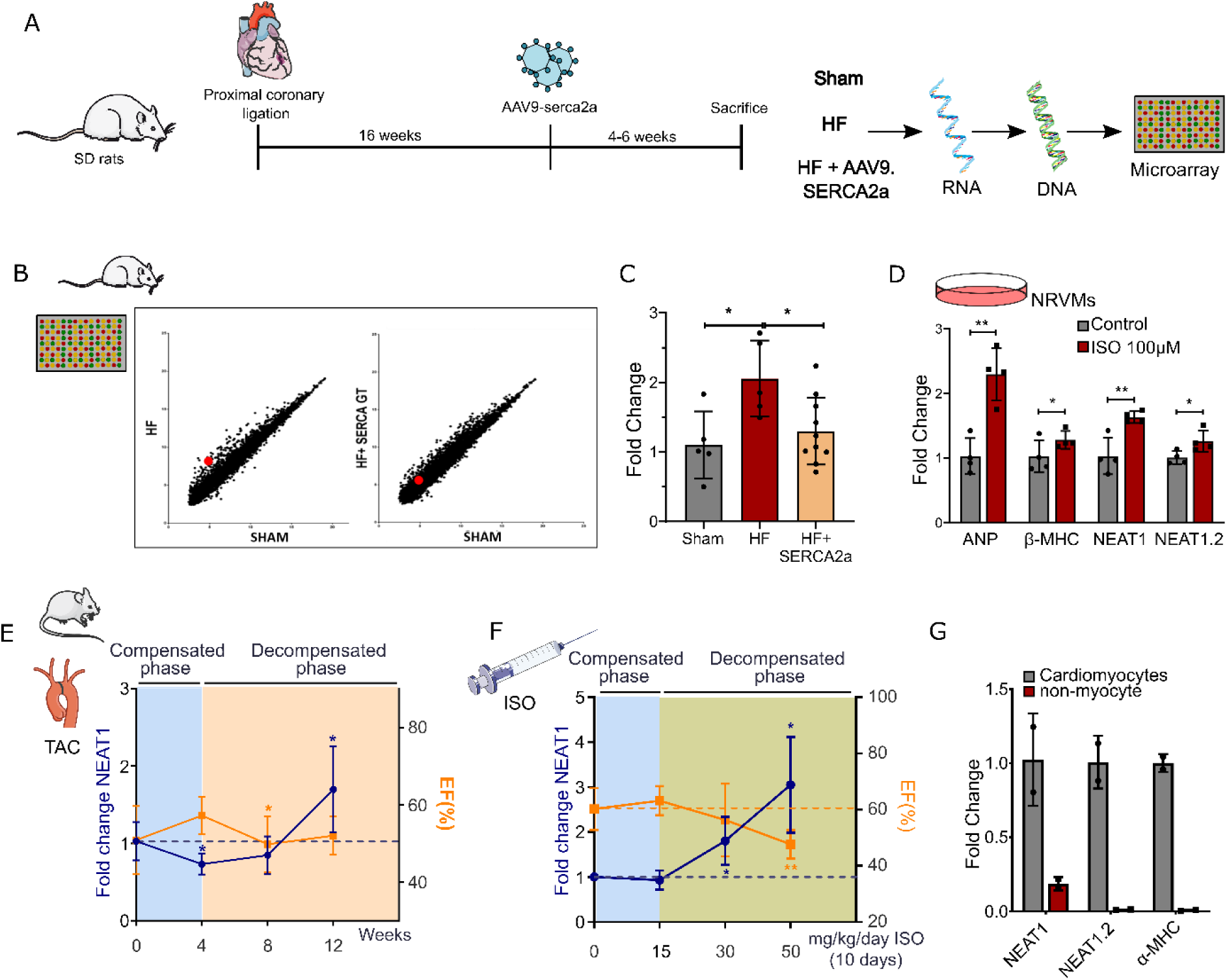
Expression of Neat1 increases during cardiac hypertrophy. **(A)** Experimental setup to detect the deregulated lncRNA in hearts subjected to HF, HF+AAV9-SERCA2a relative to Sham hearts **(B)** Microarray based profiling of RNA isolated from rat hearts subjected to Myocardial infarction and rescue by Serca2a gene therapy. **(C)** qRT-PCR of Neat1 from RNA isolated from rat ventricle after MI and MI+Serca2a gene therapy. **(D)** qRT-PCR for detecting the change in expression of ANP, β-MHC, Neat1, and Neat1.2 in primary neonatal rat ventricular myocytes (NRCM) challenged with 100 µM Isoproterenol for 48 h. * *p* value < 0.05, ** *p* value <0.01. t-tests were performed to compute statistical differences between both groups. **(E)** Neat1 expression levels detected by qRT-PCR and ejection fraction (EF%) throughout heart failure progression (TAC, transverse aortic constriction; *n* = 5–8). * *p* value < 0.05, t-tests were performed to compute statistical differences between Sham and TAC. **(F)** Neat1 expression levels and ejection fraction (EF%) throughout heart failure progression after subjecting mice to 15 mg/kg/day, 30mg/kg/day, and 50 mg/kg/day of isoproterenol for 10 days. * *p* value < 0.05, ** *p* value <0.01. t-tests were performed to compute statistical differences between saline and isoproterenol groups. **(G)** qRT-PCR for detecting the expression of total *Neat1* and Neat1.2 in adult mice cardiomyocytes and adult mice non-myocyte fraction.

### NEAT1 expression is increased in cardiomyopathies

We assessed the expression levels of *NEAT1* and *NEAT1.2* in cardiomyocytes derived from iPSC- of two Indian HCM patients, respectively (Figure 2A). Expression of Neat1 isoforms was elevated in both the models (Fig. 2B, C). Similarly, Neat1 transcript levels were elevated in the hearts of HCM associated transgenic mice (Fig. 2D).

**Figure 2.**
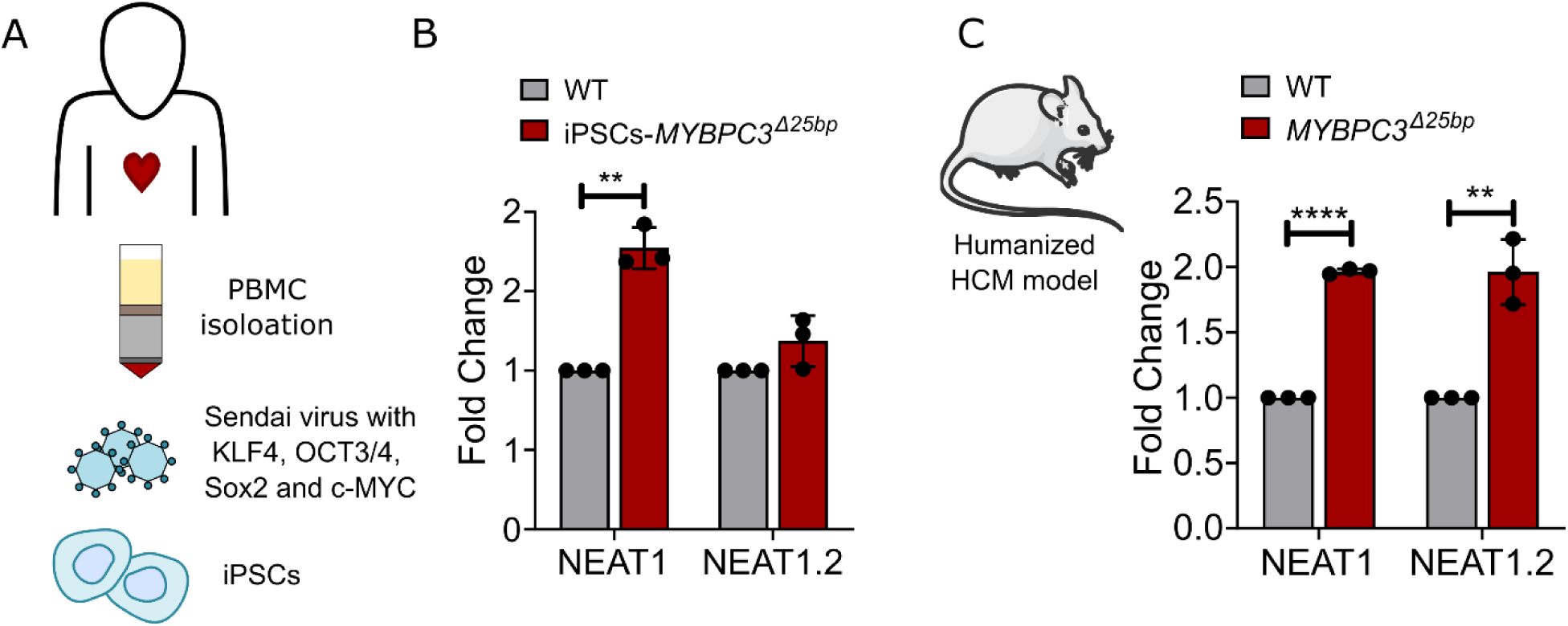
NEAT1 expression is increased in cardiomyopathies. **(A)** Schematic depicting the iPSCs preparation from patients having *MYBPC3^Δ25bp^*mutation. **(B)** qRT-PCR of NEAT1 and NEAT1.2 in cardiomyocytes derived from iPSCs hypertrophic patients having *MYBPC3^Δ25bp^* mutation. Values are shown as means ± SEM with each experiment performed in triplicate (n = 3). Significance was evaluated by unpaired student’s t test. \**p* < 0.05. **(D)** qRT-PCR of Neat1 and Neat1.2 in the heart tissue samples from hypertrophic transgenic mouse models harboring *MYBPC3^Δ25bp^*. Values are shown as means ± SEM with each experiment performed in triplicate (n = 3). Significance was evaluated by unpaired student’s t test. \**p* < 0.05.

### NEAT1 overexpression during heart failure is regulated by calcium signaling

Downregulation of SERCA2a is one of the hallmarks of heart failure. This results in impaired calcium handling, leading to the activation of calcineurin, which subsequently leads to dephosphorylation and nuclear localization of NFAT.^11, 13^ In the present study, treatment of AC16 cells with SERCA2a pharmacological inhibitor thapsigargin resulted in increased expression of RCAN1.4, indicating activation of NFAT signaling. This also coincided with increase in NEAT1 expression (Figure S4-A). Chelating calcium with BAPTA-AM was enough to lower NEAT1 expression even in the presence of thapsigargin (Figure 3A). These observations indicate that NEAT1 expression was induced by impaired calcium handling. Exploration of The Encyclopedia of DNA Elements (ENCODE) database for transcription factor-binding motifs in the NEAT1 promoter region revealed binding sites for NFAT within the 1.5kb upstream to NEAT1 transcription start site. To see if increased calcium levels leads to NFAT activation, we treated cells with calcium ionophore Ionomycin, which caused NFAT activation and enhanced NEAT1 expression (Figure 3B, S4-B). β-adrenergic stimulation with isoproterenol also increased the levels of RCAN1.4 and NEAT1 (S4-C). These results indicate a strong relationship between the activation of NFAT signaling and NEAT1 expression. Expression of a constitutively active form of NFAT (CA-NFAT2) increased NEAT1 expression (Figure 3C). Promoter luciferase reporter assay using rat NEAT1 promoter sequence in CA-NFAT2 expressing cells confirmed the regulation of NEAT1 promoter by NFAT (Figure 3D). Multiple sequence alignment revealed three conserved NFAT binding sites (TGGAA) upstream of the NEAT1 TSS (S4-D). CA-NFAT2 failed to induce NEAT1 expression when these binding sites were mutated (Figure 3D). To confirm the binding of NFAT to the NEAT1 promoter, a stable cell line having constitutive expression of EGFP-tagged NFAT was prepared. When stimulated with ionomycin or thapsigargin, the cells showed NFAT translocation in to the nucleus, which otherwise remained in the cytoplasm (Figure 3E and S4-E). The binding of NFAT on the NEAT1 promoter in these was confirmed using ChIP-qPCR. Promoter of RCAN1.4 served as positive control for NFAT binding (Figure 3F).

**Figure 3.**
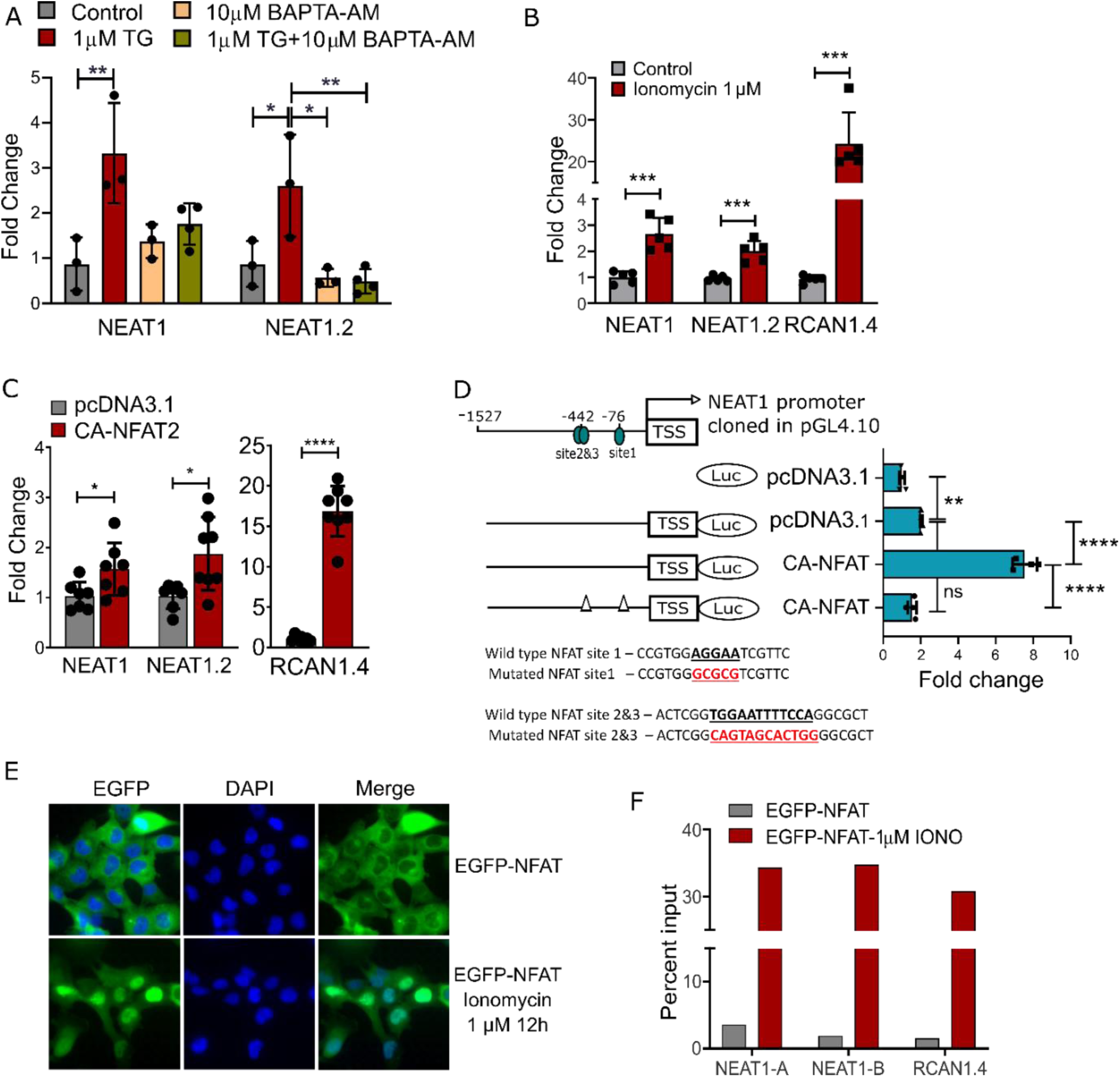
NEAT1 expression is induced by the activation of NFAT signaling and its binding on the NEAT1 promoter. **(A)** qRT-PCR of NEAT1and NEAT1.2 upon subjecting H9C2 cells with 1 μM thapsigargin, 10 µM BAPTA-AM, 1 µM thapsigargin + 10 µM BAPTA-AM. DMSO was used as the vehicle control. * *p* value < 0.05, ** *p* value< 0.01, *** *p* value <0.001. One way ANOVA was performed in multiple comparisons followed by Tukey tests. **(B)** qRT-PCR to detect changes in NEAT1, NEAT1.2 and NFAT target gene RCAN1.4 after H9C2 cells were treated with 1 µM ionomycin for 8 h. *** *p* value <0.001. t-tests were performed to compare the expression level between DMSO treated control and ionomycin treated group. **(C)** qRT-PCR to detect changes in NEAT1 expression in H9C2 cells transfected with active form of NFAT2 (CA-NFAT2). Rcan1.4 was used as a positive control for transfections. * *p* value < 0.05, *** *p* value <0.001. t-tests were performed to compare the expression level between untransfected and CA-NFAT2 transfected H9C2 cells. **(D)** Promoter luciferase assay using rat Neat1 promoter in the presence or absence of CA-NFAT and NFAT binding site on the promoter. ** *p* value< 0.01, **** *p* value <0.0001. One way ANOVA was performed in multiple comparisons followed by Tukey tests. **(E)** AC16 cells stably expressing EGFP-NFAT stimulated with 1 µM ionomycin for 12 h. An equal volume of DMSO was added as a control. **(F)** ChIP-qPCR assay to detect the NFAT binding on NEAT1 promoter using AC16 cells having a stable expression of EGFP-NFAT. Cells were treated with 1 µM ionomycin for 12h. DMSO treated cells were taken as control.

### NEAT1 is a pro-hypertrophic lncRNA

Increased NEAT1 expression is positively correlated with heart failure (Figure 1E). NEAT1 over expression (NEAT1-OE) in human cardiomyocyte cell line AC16 using NEAT1 specific gRNA (S5-A) increased the expression of ANP in the absence of any other treatment (Figure 4A). Additionally, NEAT1-OE also resulted in an increase in cell size (Figure 4B). Similar results were obtained in mouse cardiomyocyte cell line (HL-1) (S5 B, C). siRNA-based silencing of NEAT1 blunted isoproterenol induced ANP expression (Figure 4C).

**Figure 4.**
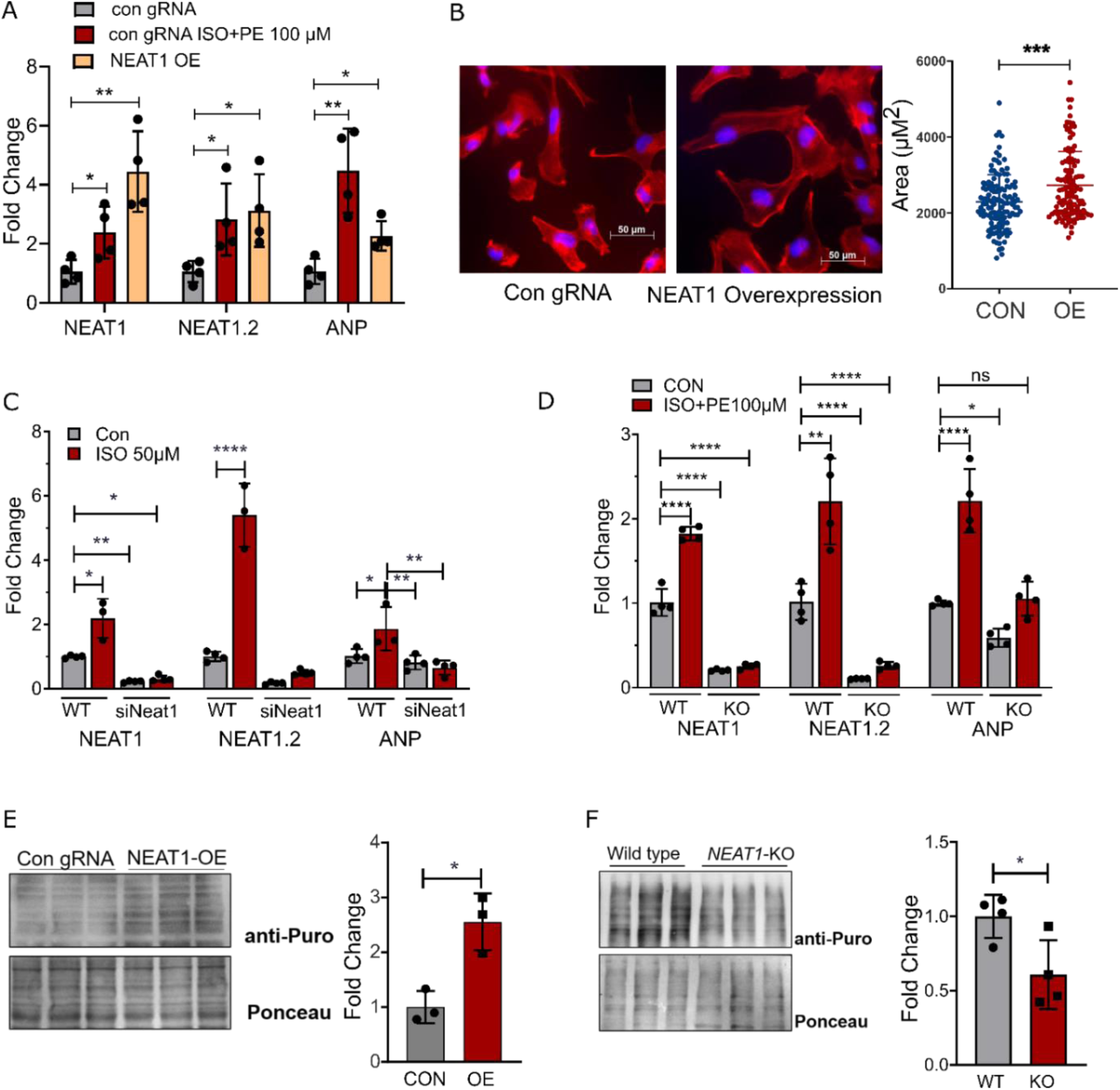
NEAT1 is a pro-hypertrophic lncRNA. **(A)** qRT-PCR to detect the changes in the level of ANP, NEAT1, and NEAT.2 expression in control AC16 cells and control cells treated with Isoproterenol and phenylephrine (100 µM each) and *Neat1-OE* cells. * *p* value < 0.05, ** *p* value< 0.01, group wise comparison with con gRNA using t-test. **(B)** Phalloidin staining of AC16 cells having *NEAT1* overexpression and control cells. Quantitative data are shown as the dot plot. *** *p* value < 0.001, t-tests were performed to compare the cell sizes. **(C)** qRT-PCR of Neat1, Neat1.2, and ANP in H9C2 cells transfected with siRNA targeting *Neat1* in the presence and absence of Isoproterenol. * *p* value < 0.05, ** *p* value< 0.01, *** *p* value 0.001, One way ANOVA was performed with multiple comparisons followed by Tukey test. **(D)** qRT-PCR of NEAT1, NEAT1.2, and ANP in AC16 WT and NEAT1-KO cells in the presence and absence of Isoproterenol. * *p* value < 0.05, ** *p* value< 0.01, *** *p* value <0.001 One way ANOVA was performed with multiple comparisons followed by Tukey test. **(E)** Western blot of the SUnSET assay performed in control and NEAT1-OE cells. Graph represents the densitometry values normalized to ponceau stained blot. * *p* value < 0.05, t-test was performed for comparison between control and NEAT1-OE. **(F)** Western blot of the SUnSET assay performed in control and NEAT1-KO cells. Graph represents the densitometry values normalized to ponceau stained blot. * *p* value < 0.05, t-test was performed for comparison between control and NEAT1-KO.

CRISPR mediated promoter disruption of NEAT1 (NEAT1-KO) was achieved by inserting the CMV-EYFP-terminator cassette just upstream of the NEAT1 transcription start site (S6 A,B).^14^ Stimulation with β-agonists resulted in increased NEAT1 and ANP expression only in wild type cells but not in NEAT1-KO cells (Figure 4D).

During cardiac hypertrophy, increase in cardiomyocyte size is accompanied by an increase in the rate of protein synthesis.^15^ Puromycin incorporation based surface sensing of translation (SUnSET) assay revealed increased rate of protein synthesis in NEAT1 over expressing cells (Figure 4E) while NEAT1-KO cells showed an opposite effect (Figure 4F). These results indicate that NEAT1 is a pro-hypertrophic lncRNA.

### NEAT1 regulates expression of its neighboring genes

To gain more insight on genes regulated by NEAT1, RNA-sequencing was performed in NEAT1-KO cells (Figure S6-C, Table S4). Gene ontology of downregulated genes indicated an enrichment of genes associated with calcium binding and cardiac muscle contraction (Figure S6-D, Table S5). Positional gene set enrichment analysis revealed a decrease in the expression of genes present in the vicinity of the NEAT1 locus. (Figure 5A, Table S6). Stronger downregulation was observed in genes that are closer to the NEAT1 TSS (Pearson’s coefficient r = 0.44; p < 0.0001) (Figure 5B). Downregulation of NEAT1 neighboring genes was also confirmed in independent qPCR experiments using NEAT1-KO cells (Figure S7-A). NEAT1 over expression has an opposite effect on expression of these genes (Figure S7-B). NEAT1 is a nuclear lncRNA and is known to interact with several genomic regions.^16^ Superimposing the earlier reported genome-wide NEAT1 binding sites,^16^ with the downregulated genes upon NEAT1-KO in the present study revealed significant overlap. (Figure 5C), indicating that NEAT1 may regulate gene expression by interacting with the chromatin. SILAC-based quantitative mass spectrometry was performed to see whether changes in transcript levels upon NEAT11 KO translate to changes in the proteome. Similar to the RNA-sequencing results, genes adjacent to NEAT1 (on chr11q13.2) are particularly downregulated at protein level as well (Figure 5D, Table S7, S8). These results indicate that NEAT1 may regulate gene expression, especially of the genes that are in close proximity.

**Figure 5.**
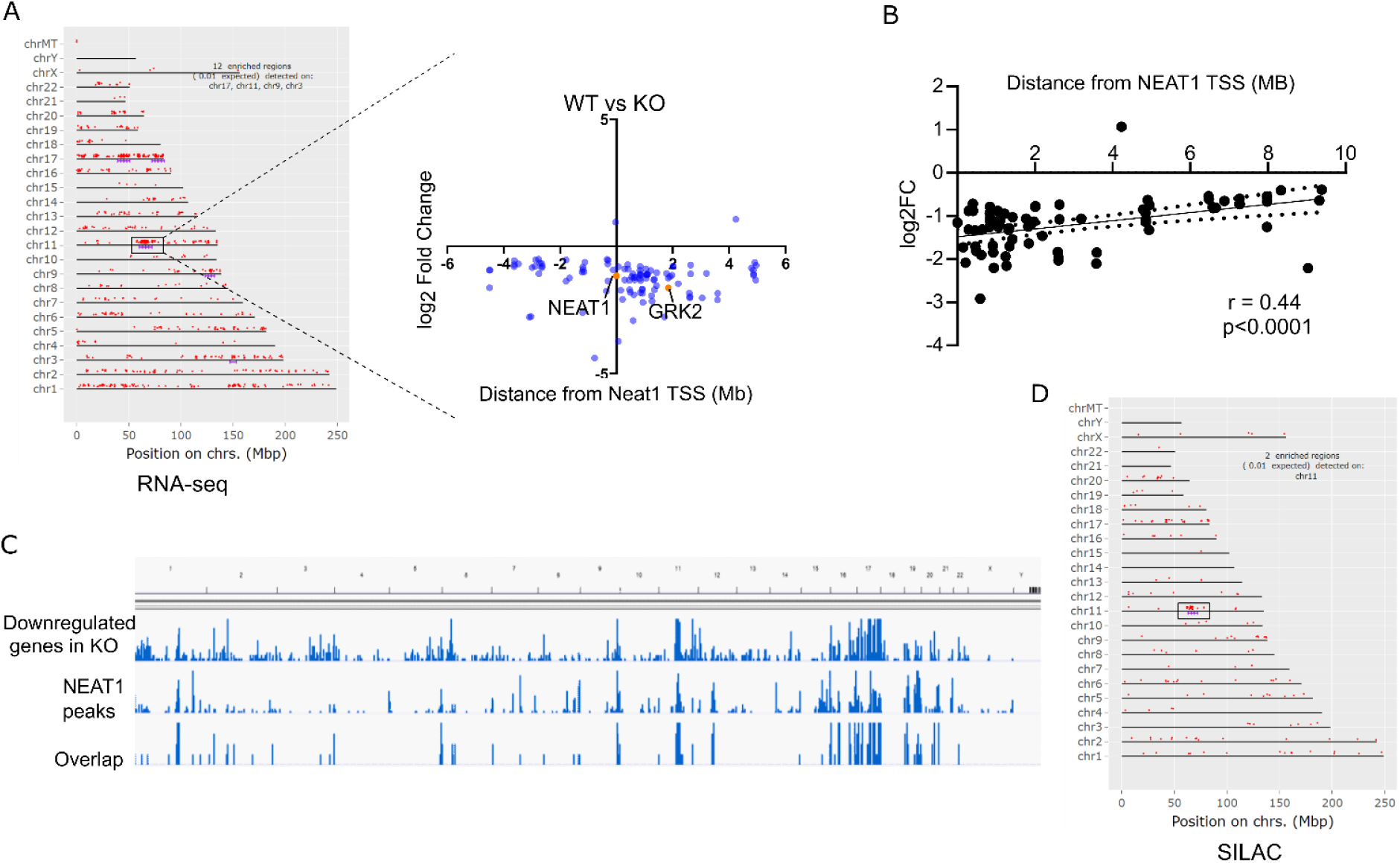
NEAT1 regulates the gene expression in cis-manner. **(A)** Chromosomal ideograms depicting the location of the genes downregulated in NEAT1-KO cells as detected by RNA-seq. The region in the black boxes indicates the enrichment of the genes in the vicinity of NEAT1. Scatter plot shows the differential expression of the genes (± 6Mb) of NEAT1 TSS between NEAT1-KO and WT cells. **(B)** Linear regression between gene expression (log2FC) and distance of genes from NEAT1 TSS. Correlation was assessed using Pearsons correation (r= 0.44, *p* value <0.0001). **(C)** qRT-PCR validation of a few NEAT1 neighboring genes in WT vs NEAT1-OE cells. **(D)** qRT-PCR validation of a few NEAT1 neighboring genes in WT vs NEAT1-KO cells. **(E)** Peaks of gene downregulated in NEAT1-KO cells, NEAT1 bound genomic regions, and overlap among both the sets. **(F)** Chromosomal ideograms depicting the location of the genes downregulated in NEAT1-KO cells as detected by SILAC. The region in the black box indicates the enrichment of the genes in the vicinity of NEAT1.

### NEAT1 interacts with p300 to control the gene expression of GRK2

G Protein-Coupled Receptor Kinase 2 (GRK2) which is also known as β-adrenergic receptor kinase is required for β-adrenergic receptor desensitization in the heart, and it has been reported that there is a 50-56% reduction in adrenergic receptor density in failing human hearts due to an increase in GRK2 expression.^17, 18^ Pharmacological inhibition of GRK2 or gene therapy using the GRK2 inhibitory peptide βARKct improved cardiac function in several preclinical models and inhibition of GRK2 in heart failure restores cAMP depletion and improves contractility.^17–19^ Interestingly, GRK2 is located within 2 Mb from NEAT TSS and GRK2 transcript and protein levels were reduced in NEAT1-KO cells (Figure 6A, B).

**Figure 6.**
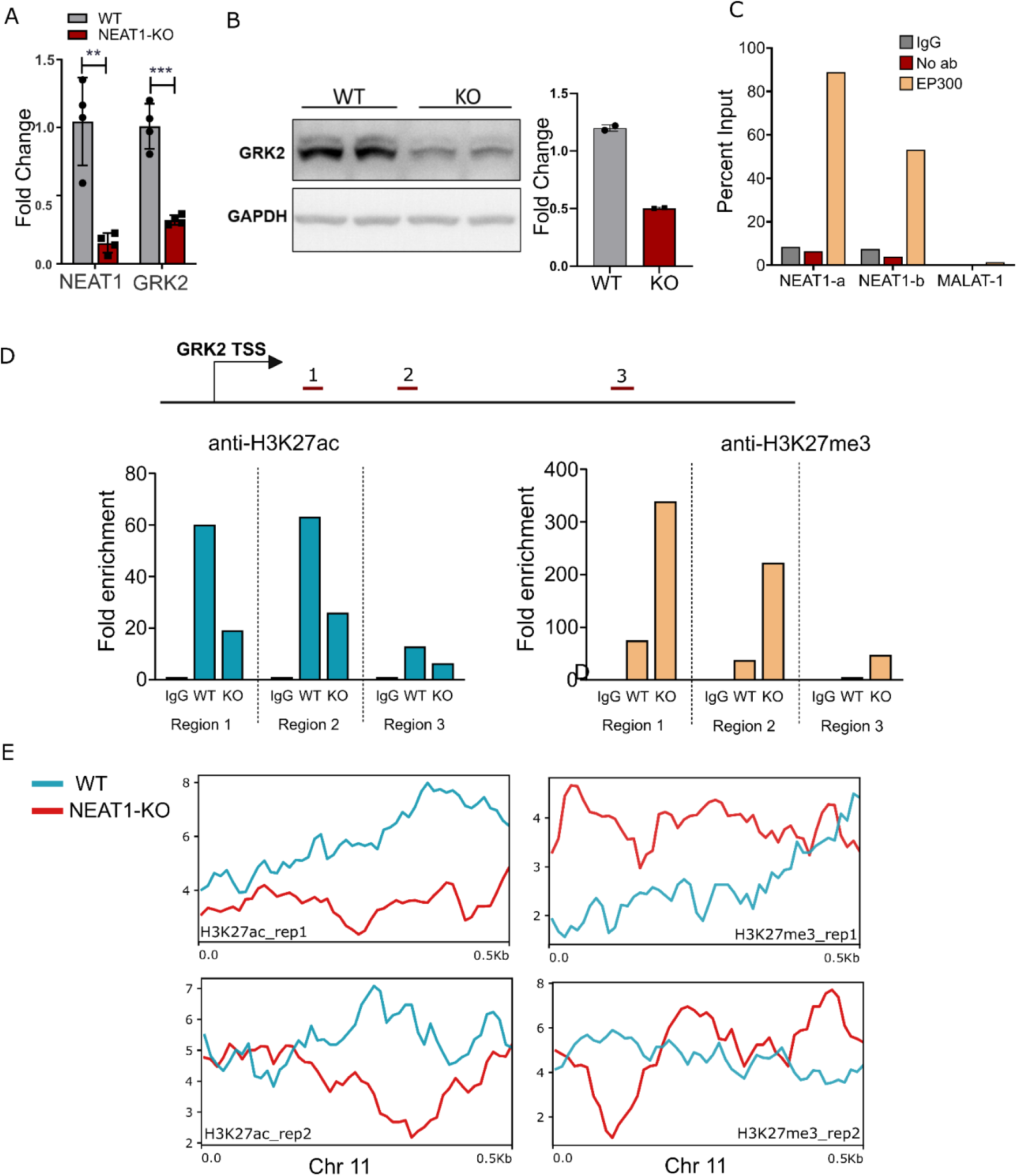
NEAT1 interacts with p300 to control the gene expression of GRK2. **(A)** qRT-PCR to measure expression of GRK2 in WT and NEAT1-KO cell lysate. ** *p* value <0.01, *** *p* value <0.001. t-tests were performed to compute statistical differences between both the groups. **(B)** Immunoblotting was performed using WT and NEAT1-KO cell lysates using anti-GRK2 and anti-GAPDH antibodies. Densitometric analysis plotted as a bar graph adjoining the blot. **(C)** RNA immunoprecipitation of lysates from AC16 cells using anti-p300 antibody and IgG as negative control. Isolated RNA was subjected to qPCR using primers specific for NEAT1, NEAT1.2, and MALAT1. **(D)** Position of primers used for assessing repressed chromatin state at GRK2 promoter in NEAT1-KO cells. Graph depicts the ChIP-qPCR from lysates of WT and NEAT1-KO cells subjected to IP using anti-H3K27ac and anti-H3K27me3 antibodies. **(E)** RPKM normalized read density plots of H3K27ac and H3K27me3 histone marks for reads aligned at Chr11 in WT and NEAT1-KO cells.

Nuclear lncRNAs often control the transcription rate by interacting with chromatin modifiers such as Brg1, PRC2, Suz12, and EZH2.^20, 21^ NEAT1, a nuclear-enriched and chromatin-associated transcript (S5 D), also serve such functions. The reduction in gene expression upon NEAT1 KO suggests that the presence of NEAT1 has an activating role at these loci. Recent reports have suggested that CBP/p300 possesses RNA-binding properties.^22^ CBP/p300 is a coactivator with histone acetyltransferase activity. A previous study reported that NEAT1 can bind to CBP/p300 to recruit the latter to promoters of the target genes.^23^ Acetylation marks on H3 lysine 27 (H3K27ac) are associated with the active promoter/enhancer elements.^24^ RNA immunoprecipitation using anti-p300 antibody indicated p300-NEAT1 interaction (Figure 6C). In line with this, chromatin immuno-precipitation using specific antibody revealed decreased H3K27 acetylation at the GRK2 promoter upon NEAT1 knock-down. Supporting these results, H3K27me3 marks on GRK2 promoter (which are marks of gene repression) were highly enriched in NEAT1 KO cells (Figure 6D).

To get a global picture, we profiled the changes in the genome wide binding of H3K27ac and H3K27me3 in WT vs KO cells using CUT&RUN protein-DNA interaction profiling. Hierarchical clustering of the reads obtained from both the cell types showed significant difference in the occupancy of H3K27ac and H3K27me3, however, both WT and KO were clustered together indicating no major differences in the occupancy between both cell types (Figure S7). Using the reads mapped to Chr11, we observed decreased occupancy of H3K27ac and increased occupancy of H3K27me3 in KO cells (Figure 6E). Overall, these results indicate that NEAT1 is acting in a cis-manner to regulate neighboring gene expression by recruiting CBP/p300.

### *Neat1^-/-^* mice can tolerate isoproterenol insult

To test the role of Neat1 in *in-vivo* settings, we subjected *Neat1^-/-^* mice and WT littermates to isoproterenol insult (60mg/kg/day) for 15 days (Figure 7A). There was no difference observed in heart weight of the untreated WT and KO mice. However, the heart weight of KO mice did not show any increase even after administrating isoproterenol, unlike the heart of WT mice which exhibited significant increase in heart weight (Figure 7B). Additionally, only WT but not KO mice exhibited an increase in cardiomyocyte size upon isoproterenol treatment (Figure 7C). Similarly, while WT mice displayed increased fibrosis upon isoproterenol stimulation, there no significant difference in fibrosis in KO mice treated with saline or isoproterenol (Figure 7D).

**Figure 7.**
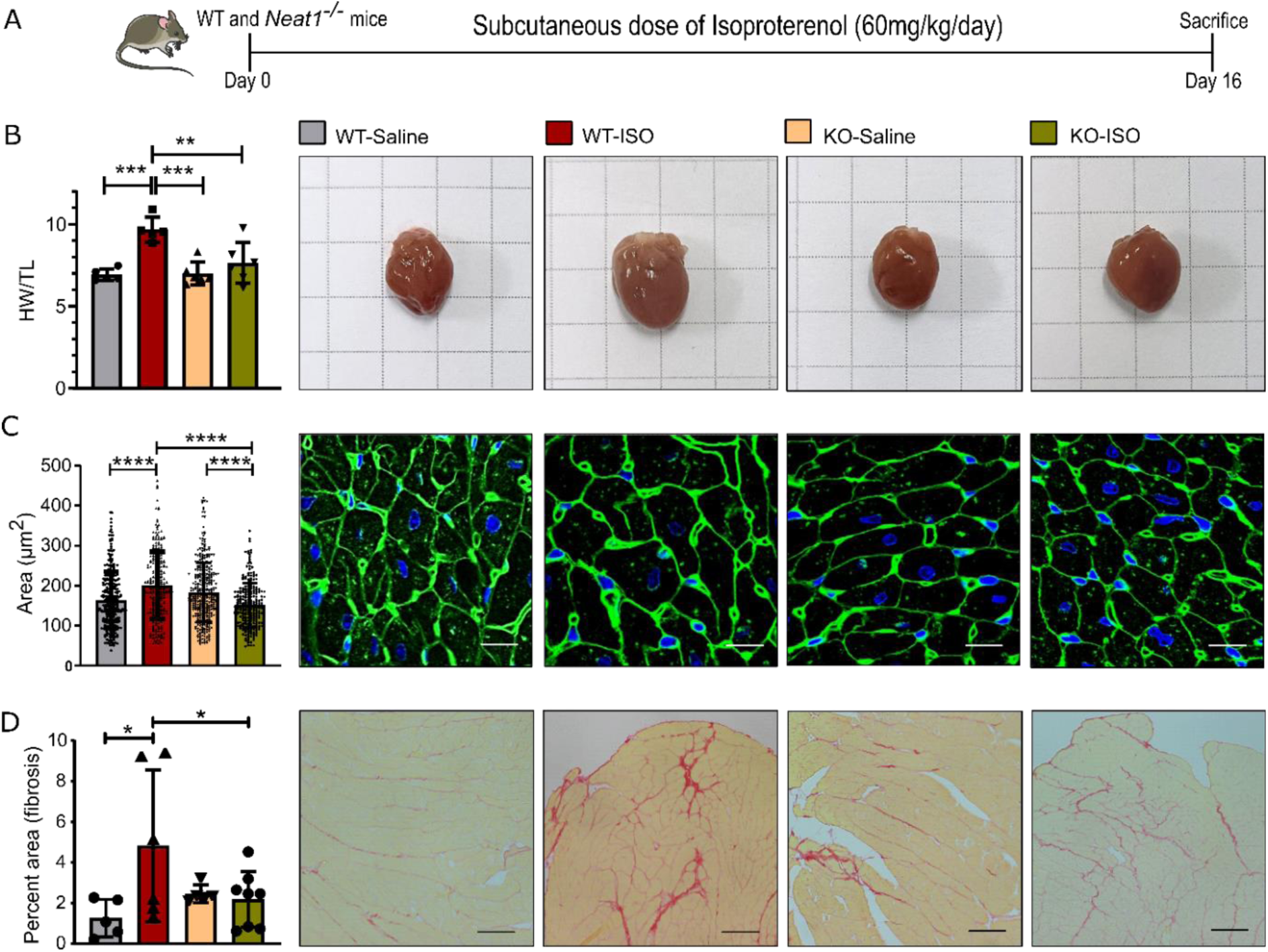
*Neat1*^-/-^ can tolerate isoproterenol insult. **(A)** Schematic representation of the experimental design. **(B)** Heart weight-to-tibia-length ratio (HW/TL) of *Neat1^-/-^*and wild-type littermates (WT). * *p* value < 0.05, ** *p* value< 0.01, *** *p* value <0.001. One way ANOVA was performed in multiple comparisons followed by Tukey tests. **(C)** Cardiomyocyte size measured after WGA staining (n = 4; DAPI = 4′,6-diamidino-2-phenylindole; WGA, wheat germ agglutinin coupled to Alexa Flour 488) of the cross-section of heart. * *p* value < 0.05, ** *p* value< 0.01, *** *p* value <0.001. One way ANOVA was performed in multiple comparisons followed by Tukey tests. **(D)** Percent area of fibrosis as after picro-sirius red staining of the cross-section of heart. * *p* value < 0.05, ** *p* value< 0.01, *** *p* value <0.001. One way ANOVA was performed in multiple comparisons followed by Tukey tests.

## Discussion

Recent studies have emphasized the role of lncRNAs in the pathophysiology of the heart. In this study, we found that lncRNA NEAT1 is upregulated in cardiac hypertrophy and operates in a cis-manner to maintain the expression of GRK2/bark1 by recruiting co-activator p300 at its promoter. Our results are consistent with previous studies showing increased NEAT1 expression post-myocardial infarction in rat hearts.^25^ Ge et al also reported an increase in NEAT1 expression in TAC mice and they further show that overexpressing NEAT1 elevates cardiac fibrosis by suppressing Smad7 transcription via Ezh2.^26^ Similarly, we observed an increase in Neat1 expression in failing left ventricle tissue from mice that were genetically, (Figure 2) surgically, (Figure 1E) or pharmacologically (Figure 1F) intervened to induce cardiac hypertrophy. NEAT1 over-expression in was sufficient to induce hypertrophic growth in cardiomyocytes and its inhibition prevents isoproterenol induced cardiomyocyte hypertrophy (Figure 4).

ENCODE data has revealed multiple transcription factor binding sites on the NEAT1 promoter. Previous studies have shown that NEAT1 is regulated by various transcription factors, such as p53,^27^ ERα,^28^ ATF2^29^ and HIF2α^30^. In this study, we have identified NFAT as a positive regulator of NEAT1 expression. NFAT signaling is known to be highly active during pathological hypertrophy.^13^ Increasing cytosolic calcium levels by ionomycin, thapsigargin, or β-agonist stimulation results in elevated NEAT1 levels (Figure 3). NFAT binding on the NEAT1 promoter may partially contribute to its increase during cardiac hypertrophy, indicating that NEAT1 is sensitive to changes in intracellular calcium levels.

RNA-sequencing experiments using NEAT1-KO cell lines revealed that genes around NEAT1 locus are particularly down regulated in the absence of NEAT1 (Figure 5). NEAT1 forms paraspeckles, which are phase-separated structures, and previous research has shown that the first paraspeckle foci appear near the NEAT1 transcription start site on Chr11 after mitosis.^31^ Previous studies have shown that NEAT1 associates with several transcriptional repressors and coactivators such as Suz12, Ezh2,^32^ CARM1,^33^ ASXL1,^34^ and ISWI^35^. Transcriptomics data combined with the genome-wide binding analysis of NEAT1, suggest its role in promoting transcription, potentially through p300 or p300 interacting chromatin remodelers. A study by Alver et al. demonstrated that SWI/SNF proteins responsible for evicting the nucleosomes, are essential for the activity of p300^36^. Additionally, core paraspeckle proteins interact with SWI/SNF remodelers. Combining these findings, we can conclude that NEAT1 is necessary for maintaining open chromatin through its role in paraspeckles.^37, 38^ The decrease in active transcription at genome wide NEAT1 binding sites suggests that NEAT1/paraspeckles can function as a local hub enriched with chromatin remodelers, and co-activators and other proteins, contributing to maintenance of transcription. This highlights the complexity of NEAT1’s role in transcriptional regulation and underscores the need for further investigation.

The adrenergic nervous system plays a crucial role in heart failure, characterized by elevated levels of circulating catecholamines.^39^ Initially, this increased activity enhances ionotropy and contractility, serving as a protective mechanism. However, in later stages, the heightened workload overwhelms the cardiac muscles. To mitigate the heightened ionotropy, G-protein receptor kinase 2 (GRK2) levels are elevated in the heart. ^18, 40, 41^ This upregulation of GRK2 has been observed in both human heart failure patients and various preclinical models of heart failure. GRK2, also known as β-adrenergic receptor kinase 1 (BARK1), phosphorylates the β-adrenergic receptor, rendering it susceptible to β-arrestin-mediated receptor internalization and subsequent reduction in membrane βAR levels, leading to a decrease in ionotropy.^42^ Prolonged elevation of GRK2 attenuates βAR signaling and depletes cardiac contractile reserves. Several preclinical studies have demonstrated that reducing GRK2 activity, either through chemical inhibition using paroxetine hydrochloride or via ectopic expression of GRK2 inhibitory peptides like βARKrgs and βARKct, can reduce cardiac hypertrophy and mitigate heart failure.^18, 43–45^

In our study, we discovered that the depletion of NEAT1 leads to a reduction in GRK2 levels (Figure 6). In summary, during cardiac hypertrophy, the increase in NEAT1 expression plays a role in upregulating GRK2 by recruiting p300 to its promoter. These finding suggests that NEAT1, at least in part, contributes to the elevation of GRK2 levels associated with cardiomyocyte hypertrophy.

## Material and methods

### Microarray

Rat lncRNA microarray (4X44K, Arraystar) was performed in the heart samples isolated from the rats that were subjected to proximal coronary artery ligation (HF), HF+ SERCA2A gene therapy. These animals were described elsewhere^12, 46^.

### Animal experiments

Animal experiments were performed according to local and national ethics guidelines. Cardiac pressure overload was generated in C57BL6/J mice (8 to 12 weeks old) using transverse aortic constriction. Using a minimally invasive technique a titanium clip calibrated around a 28 G needle was placed on the aortic arch, without opening the thoracic cavity as described previously.^47, 48^ Sham animals underwent the same surgery without placement of the clip. Clip placement was confirmed one week after surgery using color doppler (Vevo2100, VisualSonics, MS400 transducer). Animals were continuously monitored for various cardiac parameters using echocardiography. Animals were sacrificed after 4, 8, and, 12 weeks post-surgery. For pharmacologically induced hypertrophy 10 weeks old C57BL6/J male mice received daily indicated doses (15mg/kg/day, 30mg/kg/day, and 50mg/kg/day) of Isoproterenol (Tocris, #1747) subcutaneously for 10 days. Control animals were given subcutaneous injections of saline. Animals were sacrificed and their body weight, heart weight, and tibia length were measured to assess hypertrophy. Mice were euthanised and hearts were perfused with 30 mM KCl to induce diastolic arrest followed by perfusion with PBS, and the apex portion of the heart was snap-frozen in liquid nitrogen for RNA isolation, the middle cross-section was fixed in 4% neutral buffered formalin for histology, and the top section was collected for protein lysate preparation.

Animals were continuously monitored using echocardiography. Left ventricle (LV) was observed using a parasternal long-axis view (PLAX), and images of the heart were captured in both B-mode and M-mode. To calculate various cardiac parameters, LV tracing was done on the captured M-mode image. Per image, three random regions were selected for tracing the LV wall and hemodynamic parameters using the VevoLab software.

### Maintenance of *Neat1^-/-^* mice

*Neat1^-/-^* mice were procured from RIKEN, Japan ^49^. Male *Neat1^-/-^* mice were crossed with the female C57BL6/N mice and the heterozygote progeny were used for breeding. Genotyping was performed to segregate the mice and 8-12 weeks of *Neat1^-/-^* mice and wt littermates were used for the experiment. Mice were subjected to subcutaneous injections of isoproterenol (60mg/kg/day) for 15 days after which they were sacrificed as per the institute’s animal ethics guidelines.

### Transgenic humanized HCM mouse model

Previously, we showed that a 25 bp deletion in the MYBPC3(*MYBPC3*^Δ25bp^) gene affecting the C10 domain of the MYBPC3 protein is associated with cardiomyopathy. 48 To obtain an HCM model, a cardiac-specific transgenic mouse model overexpressing ∼ 50-60% of MYBPC3 mutant proteins with a modified C10 domain was generated (Tg). The Tg mice developed cardiac hypertrophy at 5 months and recapitulated HCM patient phenotypes, including an increase in heart weight to body weight ratio, displayed significant elevations in the ejection fraction (EF), fractional shortening (FS), and expression of fetal genes (Nppa and Nppb) and calcium handling gene ratios (Pln/Serca2a) compared to non-transgenic controls. These data suggest that the transgenic mouse model displayed features of HCM ^50^.

### CM derivation from induced pluripotent stem cell (iPSC) lines of two HCM patients

The iPSC lines were generated using peripheral blood mononuclear cells (PBMC) from an Indian HCM patient having *MYBPC3*^Δ25bp^ mutation ^51^. PBMCs were separated from whole blood using histopaque 1077 (Sigma) and SepMate-15 tube (StemCell technologies) by density gradient centrifugation. After four days, PBMCs were reprogrammed into iPSC by transducing CytoTune-iPS 2.0 Sendai reprogramming vectors (Thermo Fisher Scientific) expressing human KLFf4, OCT3/4, SOX2 and c-MYC, following the manufacturer’s instruction. After 18 days post-transduction, iPSC colonies were transferred to vitronectin (Thermo Fisher Scientific) coated 12 well plates and cultured using StemFlex medium (Thermo Fisher Scientific) at 37 ◦C and 5 % CO2 in a humidified atmosphere.

### Differentiation of iPSCs into cardiomyocytes

For cardiomyocyte differentiation, iPSC cells were cultured up to 80 to 90 % confluency. Differentiation was induced by treating the cells with 5µM/ml CHIR99021 (Sigma-Aldrich) for 2 days and followed by 5µM IWP2 (Sigma-Aldrich) treatment for another 2 days (freshly add up for 24 hrs once) in cardiac differentiation media (RPMI1640 supplemented with glutamax, 0.5mg/ml human recombinant albumin and 0.2 mg/ml L ascorbic acid). On day 8, cardiac differentiation media was switched to cardiac maintenance media (RPMI1640 supplemented with 2% B27 minus insulin and glutamax). Cells were maintained in this media and changed every 3 days thereafter. The cardiomyocytes beating was observed on 12th day.

### Cardiac cell fractionation

Adult mouse cardiomyocytes were isolated using the Langendorff-free method where the aorta is clamped and the heart is continuously perfused through the left ventricle with the help of a syringe.^52^ To separate the rod-shaped myocytes from non-myocytes we briefly centrifuged the cell suspension at 0.1g for 20 sec in the settling of round cardiomyocytes. All of the non-myocyte cell suspension was pooled by centrifugation at 0.8g for 5 mins and 1 ml trizol was added to both myocyte and non-myocyte cell pellet. Purity of cell population was assessed by qRT-PCR of α-MHC (primers listed in table S9).

### Neonatal Rat cardiac myocytes isolation

For the preparation of neonatal rat cardiomyocytes we used serial trypsin digestion as described previously.^53^ After 1h of preplating, the cells were cultured in MEM 5% FBS having 10mM BrDU to stop fibroblast from proliferation. For inducing hypertrophy FBS concentration was reduced to 0.5%. Cells were starved in MEM-0.5% FBS for 24 h and treated with 100μM ISO and 100μM PE for 48 h.

### Cell culture

H9C2 cells were purchased from ATCC grown in DMEM media with 10% FBS. For hypertrophy experiments, cells were starved for 18h in 0.1% FBS containing DMEM media prior to the addition of 50μM Isoproterenol (Tocris, #1747) for 24h. Cells were treated with 1μM Thapsigargin (Tocris, #1138) and 10μM BAPTA-AM (Tocris, #2787) for 8h in DMEM 10% FBS, and DMSO was used as control. HL-1 cell line was purchased from Millipore maintained in Claycomb media (#51800C, Millipore) with the addition of L-2mM L-glutamine, 10mM Norepinephrine, and 10% FBS. For hypertrophy experiments cells were starved in Claycomb media lacking FBS and norepinephrine. Cells were starved for 24 h after which 100μM Isoproterenol and 100μM Phenylephrine (Tocris, #2838) were added to the starvation media for 48h. The AC16 cell line was purchased from Millipore, grown in DMEM/F12 media containing 12.5% FBS. For hypertrophy experiments cells were serum starved for 24 h and treated with 100μM ISO+100μM PE for another 24h. Cells were treated with 1μM Thapsigargin (Tocris, #1138) for 12 h. For assessing calcium-dependent NEAT1 expression H9C2 and AC16 cells were treated with 1μM Ionomycin (Sigma Aldrich, #10634-1MG) for 8h and 12 h respectively. HEK293T cells were cultured in DMEM media with 10% FBS. All the cell lines were grown in a 5% CO_2_ incubator at 37 °C and were regularly monitored for mycoplasma.

### RNA extraction

Cells were collected in Trizol (RNAiso Plus Takara) and RNA was isolated according to the manufacturer’s instructions. Before cDNA synthesis RNA was subjected to DNaseI digestion to remove residual genomic DNA. RNA integrity was checked on an agarose gel. Samples having degraded RNA were discarded. 1ug of RNA was used for preparing cDNA using the Biorad iscript select cDNA synthesis kit according to the manufacturer’s instructions.

### Preparation of NEAT1 KO cell line

For the preparation of the NEAT1-KO cell line we used the strategy used by Ruhohan Li *et.al.*^14^ AC16 cells seeded in a 6 well, were transfected using Lipofectamine 3000 as per manufacturer’s instructions with plasmids having Cas9 and gRNA (Addgene #97081) and a plasmid having repair template (Addgene #97088) These plasmids were a gift from Archa Fox. Transfected cells were expanded to a T-75 after which cells underwent three rounds of FACS sorting and expansion to enrich the high EYFP-expressing population. Individual EYFP-positive cells were seeded in a 96-well plate for clonal expansion. After 3-4 weeks DNA was isolated and cells were screened for insertion using primers given in Table S9. Clones homozygous for insertion were tested for NEAT1 downregulation using primers given in Table S9. Transfected cells that did not undergo targeted editing were used as WT controls for subsequent experiments.

### Lentiviral production

All lentiviral work was done in the BSL-2 facility, according to the guidelines set by the institutional biosafety committee. About 3 million HEK293T cells were seeded in 100 mm dish 24 hours prior to transfection. Cells were transfected with transfer plasmid, viral packaging plasmid (pspax, Addgene #12260), and viral envelope plasmid (pMD2.G, Addgene #12259) in 4:3:1 molar ratio. Transfection was performed with Lipofectamine 3000 according to the manufacturer’s instructions. Transfection media was replaced, 4h post-transfection with 10ml of DMEM having 10% FBS. Media collected 72h post transfections and centrifuged at 700g for 10 mins. Viral particles were precipitated by adding Lenti-X Concentrator (Takara, #631232) according to the manufacturer’s instructions

### Preparation of NEAT1-OE cell lines

We used CRISPR/Cas9 Synergistic Activation Mediator (SAM) strategy to induce upregulation of NEAT1.^54^ Lentiviral particles were generated from Addgene plasmid #61425 having dCAS9-VP64 and plasmid #61426 containing MS2-P65-HSF. AC16 and HL-1 cells were transduced with concentrated viral particles in the presence of 8 μg/ml of Polybrene. Post-transduction cells were selected with hygromycin and blasticidin. AC16 and HL-1 cells were selected with 5 μg/ml of Blasticidin and 300 μg/ml of Hygromycin. For AC16 immortalized human cardiomyocytes cells we selected gRNA based on published results.^55^ For HL-1 gRNA were designed using the Broad institute GPP web portal. gRNAs were cloned into lenti sgRNA(MS2)_zeo (Addgene #61427) backbone using BsmBI restriction enzyme site. As a control, a published non-targeting gRNA was also cloned into the same backbone.^56^ Lentiviral particles were generated and cells stably expressing dcas9-VP64 and MS2-P65-HSF were transduced with concentrated viral particles in the presence of 8 μg/ml of polybrene. AC16 cells were selected using 400 μg/ml of Zeocin. Cells were selected for about 14 days. Upregulation of NEAT1 was confirmed using qRT-PCR. This upregulation was compared to the cells transduced with non-targeting gRNA sequences. Induction of hypertrophy was done as described above.

### Promoter Analysis and Luciferase Assay

Genomic DNA from rat tails was used for promoter amplification. The amplicon was inserted into pGL4.10 after digestion with KpnI and HindIII. The constitutively active form of NFAT2 was cloned from pENTR11 CA NFAT2 (Addgene #11793) into pEZY3 plasmid (Addgene #18672) using Gateway cloning. pEZY3 plasmid is a gateway-compatible plasmid derived from pcDNA, hence, pcDNA3.1 was used as an empty control. NFAT mutagenesis was performed using overlap extension PCR using primers given in table S9

For performing the luciferase assay, we used NIH3T3 cells. Cells were transfected with NEAT1 promoter constructs with or without CA-NFAT2. Renilla-Luc (pRL-SV40) was used as an internal control. 24h post-transfection luciferase activity was measured using a Dual luciferase assay kit (#E1910) from Promega according to the manufacturer’s instructions. Fold change was calculated with respect to cells transfected with NEAT1 promoter and pcDNA3.1.

### Phalloidin staining

Cells seeded on a coverslip were fixed with 4% formaldehyde in PBS for 10 min at 37℃. After permeabilizing with 0.1% TritonX in PBS for 15 mins, 6 μM of Alexa Fluor™ 594 Phalloidin was prepared in 1% BSA-PBS and added to each coverslip and kept in the dark for 40 minutes after which cells with PBS and coverslips were mounted on the slides with vectashield mountant containing DAPI. At Least five images were taken per coverslip using Axioimager Z and cell size was measured using Axio-vision Rel. 4.8 software. Cell size measurement between WT and Neat1-OE cells was done in a blind manner.

### Histology / WGA staining

Tissue cross-sections collected from the heart were dehydrated by passing them through a series of increasing ethanol concentrations followed by paraffin embedding. Embedded tissues were sliced into sections of 4 microns and placed on charged slides. Sections were deparaffinized and rehydrated. Wheat Germ Agglutinin, Alexa Fluor™ 488 Conjugate (#W11261, Invitrogen) was diluted 1:100 from a stock solution of 1 mg/ml and placed over each section. Sections were kept in the dark for 40 mins followed by PBS wash and counterstained with DAPI 1 μg/ml dissolved in PBS. Slides were washed thrice with PBS and the coverslip was mounted using Prolong Gold antifade medium. Images were taken by Olympus FV3000 confocal microscope at 60X. Two images were taken per coverslip. Cell size was measured using ImageJ, all the myocytes per section were scored.

### ChIP

2 X 10^6 cells were collected per replicate. These cells were fixed with 1% methanol-free formaldehyde for 20 min. Cell pellets were thawed at 4℃ and resuspended in 200μl of ChIP lysis buffer (1%SDS, 10 mM EDTA, 50 mM Tris, pH 8.1) and transferred in a Bioruptor tube (#C30010016.) for sonication. Sonication was done at 4 ℃ for 8 cycles of 20 seconds ON, 30 seconds OFF. The lysate was cleared of any debris by spinning at 12000 g for 10 mins and the clear supernatant was transferred to a clean tube. 70 μg chromatin was taken per IP and volume was made up to 1 ml by adding ChIP dilution buffer (0.01% SDS, 1.1% Triton X-100, 1.2mM EDTA, 16.7mM Tris-HCl, pH 8.1, 167mM NaCl) and 100 μl was taken out from each tube as INPUT. For assessing histone modification 70 μg of chromatin from WT and KO cells was taken and 3ug antibodies were added H3K27ac (Active motif, #39134), H3K27me3 (Abcam, ab195477). This was kept for overnight incubation at 4℃ on an end-to-end rotor. 50 μl of protein A beads/sample were washed and added to the lysates and kept at 4℃ for 2 h. The beads were collected and washes were repeated with low salt buffer (0.1% SDS, 1% Triton X-100, 2 mM EDTA, 20 mM Tris-HCl, pH 8.1, 150 mM NaCl), followed by high salt buffer (0.1% SDS, 1% Triton X-100, 2 mM EDTA, 20 mM Tris-HCl, pH 8.1, 500 mM NaCl), LiCl buffer (0.25 M LiCl, 1% IGEPAL-CA630, 1% deoxycholic acid (sodium salt), 1 mM EDTA, 10 mM Tris, pH 8.1) and 2 washes with TE buffer (10 mM Tris-HCl, 1 mM EDTA, pH 8.0). 200μl of freshly prepared 1% SDS in NaHCO3 prewarmed to 65℃ to the beads was added and vortexed. This supernatant was collected and elution was repeated again. To 400μl of eluate and INPUT, samples 20ul of 5M NaCl and was added and eluate was kept at 55℃ overnight for reverse crosslinking. Next samples are traeted with RNAse A for 30min at 37℃ followed by ProteinaseK at 37℃ for 30min. DNA extraction was performed by Phenol-chloroform.

### Chromatin Immunoprecipitation using GFP-nanobody

GFP nanobody beads were prepared from vector pGEX6P1-GFP-Nanobody (Addgene #61838) as per the protocol.^57^ The beads were stored in 20% ethanol made in PBS as 1:1 slurry. Lentiviral mediated stable cells expressing EGFP-NFAT were stimulated with 1μM ionomycin and DMSO was used as vehicle control. After 12h of Ionomycin, 2X10^6 cells were collected per replicate. ChIP was performed as described above with few modifications. Equal amount of lysate was collected from untreated and treated cells and 20 μl of GFPnb beads were added per pulldown and incubated at 4℃ overnight. Washes and elution was done as per ChIP protocol described above.

### RNA-IP

2 x 10^6 cells (AC16 cells) were harvested and crosslinked with 1% formaldehyde in PBS for 10 min on an end-to-end rotor at RT. Antibody bead complex was prepared by resuspending 30µl washed protein G beads per pulldown using RIP buffer. The washed beads were divided equally in volume up to 100 µL and 1µg of anti-p300 (#ab14984), 1µg IgG isotype (abcam, ab171870) was added to the two tubes and one was kept as no antibody control. The tubes were incubated for 2h at room temperature. Before addition to the lysate, the antibody-bead complex was washed thoroughly at least three times with RIP buffer. Simultaneously, the fixed cell pellet was resuspended in 300µL lysis buffer (10 mM Tris-HCl, pH 7.4, 10 mM NaCl, 0.5 % NP-40, 1 mM DTT, 200 units/ml RNase inhibitor, and EDTA-free Protease Inhibitor Cocktail) and transferred to a 1.5 ml TPX tube from the diagnosis node. The lysate was sheared at 4°C using a Bioruptor Pico for 30 s ON and 30 s OFF until it became clear. The sheared lysate was cleared by centrifugation at 12,000 × g for 10 min, the cleared supernatant was divided into three tubes, and the volume was made up to 1 ml. 100 µl was removed as 10% input and antibody-bead complexes were added to the lysate. The tubes were kept on an end-to-end rotor at 4°C overnight. Beads were pulled down using a magnetic stand and washed thrice with 1 ml RIP buffer at 4°C for 5 min, followed by a PBS wash. To the beads, 200 µL of PBS and 5µl of 20 mg/ml proteinase K and SDS up to 1% were added, and the tubes were kept at 37°C for 1h. Once reverse crosslinking was performed, 800 µL of TRiZol was added and RNA was isolated. RNA was converted to cDNA and subjected to qPCR using NEAT1 specific primers. As a negative control we also tested MALAT1.

### Cell fractionation

Fragmentation of AC16 cells into cytoplasmatic, nuclear-soluble, and chromatin-associated fractions was performed as described before.^58^ Briefly, cells from a 10 cm confluent dish were collected and washed with PBS. The pellet was resuspended in 300μl of NP40-lysis buffer (10 mM Tris pH 7.4, 150 mM NaCl, 0.15 %, NP40, in RNAse-free water) and incubated on ice for 10 min. This lysate was overlayed on top of a sucrose buffer (10 mM Tris pH 7.4, 150 mM NaCl, 24 % sucrose) by pipetting to the wall of the tube. This was centrifuged at 3,500g for 10min and the supernatant (cytoplasmic fraction) was separated into a fresh tube. Cytoplasmic fraction was cleared by centrifugation at 14,000g for 1 min and to clear supernatant 800μl of trizol was added.

The pellet (nuclear fraction) was washed 3 times with ice cold PBS-EDTA by spinning at 3000 rpm for 5 min. For chromatin separation, nuclear pellet was resuspended in 250μl of glycerol buffer and 250μl of urea buffer was added immediately. Tubes were vortexed for 4s and kept on ice for 2 min. Lysate was centrifuged at 13,000g for 2 min and supernatant (nucleoplasm) was separated into a new tube and trizol was added. The pellet (chromatin fraction) was washed with PBS-EDTA and the pellet was dissolved in trizol. RNA from all fractions was isolated and converted to cDNA. Localization of NEAT1 was checked using primer in table S9. As positive control primer for pre-gapdh was used.

### SILAC

AC16 cells were grown in SILAC DMEM/F12 (Thermo Scientific) supplemented with 12.5% dialyzed fetal bovine serum (Gibco). Labeled cell lines were prepared by growing cells in suitable media for 5 passages. WT cells were grown in light [l-lysine 2HCl/l-arginine HCl (Lys0/Arg0) and NEAT1-KO AC16 cells were labeled with heavy [l-lysine 2HCl (13C6, 15N2)/l-arginine HCl (13C6, 15N4) (Lys8/Arg10)] isotopes of lysine and arginine (Thermo Scientific). All labeled cells were seeded on a P60 dish, once 70% confluent, for 24h for inducing hypertrophy. Light-labeled cells served as control. Cells were collected, and counted, 1.5x10^6 cells of each type were pooled for lysate preparation.

### Sample preparation for Mass Spectrometry

The pooled lysate was separated on NuPAGE 4%-12% bis-Tris Protein Gels (Invitrogen). Gel was run in MES buffer (100 mM Mes, 100 mM Tris–HCl, 2 mM EDTA, 7 mM SDS) at 200V for 35 min, stained with coomassie brilliant blue. Prepared gel slices underwent reduction and alkylation followed by trypsin digestion as described previously.^59^ Peptides were extracted and desalted using Pierce™ C18 Tips, 10 µL bed zip tip and eluted using 0.1% TFA and 50% ACN solution as described previously.^60^

Raw files were analyzed using MaxQuant. For peptide identification, raw MS data files were analyzed using MaxQuant,^61^ and searched against the Uniprot database of *Homo sapiens* (release 2019.03 with 42,419 entries) and a database of known contaminants. MaxQuant used a decoy version of the specified database to adjust the false discovery rates for proteins and peptides below 1%. The search parameters included constant modification of cysteine by carbamidomethylation, enzyme specificity trypsin, and multiplicity set to 2 with Lys8 and Arg10 as heavy label. Other parameters included minimum peptides for identification 1, minimum ratio count 1, and the requantify option was selected. Statistical analysis was performed in the Perseus environment.

### RNA-seq

DNaseI digested RNA extracted from the WT cells and Neat1-KO AC16 cells was sequenced in replicates of 4. Sequencing libraries were prepared according to Illumina TrueSeq total RNA kit. Quality control of raw reads was done by FastQC.^62^. Cutadapt v4.1 was used to filter the raw reads - trimmed the adapter sequence, removed the low-quality bases with a quality cutoff of 20, and discarded reads with a minimum read length of 50.^63^ Bowtie2 aligner with default parameters was used the trimmed reads to the hg38 assembly human reference genome.^64^ Principal Component Analysis (PCA) was conducted on the rlog normalized counts to eliminate outliers from the dataset. Differential gene expression analysis was performed using Deseq2.^65^ Sorted gene list from deseq2 was used to perform Gene set enrichment analysis (GSEA) using R package of cluster profiler. Bed file of NEAT1 binding peaks from West et al. was converted to hg38 coordinates using liftover. IGV was used for genome wide visualization of the peaks.

### Cut&Run

H3K27me3 and H3K27ac mapping carried out according to Cut & Run protocol. IgG used as negative control. Nuclei of ∼1×10^5^ WT and KO cells were bound to ConA magnetic beads (#BP531) and the Cut&Run protocol as described before.^66^ After targeted fragments were released, crosslink reversed by incubation at 65 °C with 2 μl 10% SDS and 2.5 μl Proteinase K (20 mg/ml) overnight. DNA was phenol/chloroform extracted and EtOH precipitated in the presence of 2 μl glycogen (2 mg/mL). Samples were recovered in 25 μl 1 mM Tris-HCl pH8, 0.1 mM EDTA. For analysis, DNA libraries were prepared according to the manufacturer’s protocol (NEBNext® Ultra II, NEB). Sequencing was performed on the NovaSeq Illumina). The resulting raw reads were assessed for quality, adapter content and duplication rates with FastQC^62^. Trim Galore was used to trim reads before alignment to the human genome hg38 using Bowtie2.^64, 67^ Duplicate reads were removed with Picard 2^68^. Deep tools was used to generate multibam summary which was used for plotHeatmap (deepTools Version 3.5.1) to generate heatmaps of H3K27ac and H3K27me3 binding across the genome. Distribution of H3K27ac and H3K27me3 reads of NEAT1 KO and WT at NEAT1 binding regions from chr11 (peaks coordinates from West et. al ^16^) were plotted as RPKM normalized read density plots using the plotProfile tool.

### Statistical analysis

GraphPad Prism 9 was used for making graphs and performing statistical analysis. All data is expressed as mean ± SD values. Significant differences are indicated as *p<0.05, **p<0.01, ***p<0.001, ****p<0.0001. two-tailed unpaired t-test was performed to compare two groups. More than two groups were analyzed using one way ANOVA. Cell size measurement done in Figure 4B and supplementary figure 5C were performed in double blind manner. Detailed information is provided in each figure legend.

## Acknowledgements

This study was supported by DBT/Wellcome Trust India Alliance (India Alliance) fellowship grant (fellowship No. IA/I/16/1/502357), Indian Council of Medical Research (grant No. small0384) and Science and Engineering Research Board (CRG/2023/007877) to RK. PP and DN received fellowship from University Grants Commission (UGC), India. PS received fellowship from Council of Scientific & Industrial Research (CSIR), India. We thank Tanya Agarwal and Divya Tej Sowpati for analyz. We thank the NGS, Proteomics, Advanced microscopy, histology, and cell culture facility at CSIR-Centre for Cellular and Molecular Biology for their assistance.

## Conflict of interest

TT is founder and shareholder of Cardior Pharmaceuticals GmbH. Others have no conflict of interest.

## Data Availability

The supplementary data derived from the RNA seq and SILAC are given as excel file along with the manuscript.

## Notes

### Competing Interest Statement

Thomas Thum is the founder and shareholder of Cardior Pharmaceuticals GmbH. Others have no conflict of interest.

